# Adaptive evolution of *Moniliophthora* PR-1 proteins towards its pathogenic lifestyle

**DOI:** 10.1101/2021.03.07.434232

**Authors:** Adrielle A. Vasconcelos, Juliana José, Paulo M. Tokimatu Filho, Antonio P. Camargo, Paulo J. P. L. Teixeira, Daniela P. T. Thomazella, Paula F. V. do Prado, Gabriel L. Fiorin, Juliana L. Costa, Antonio Figueira, Marcelo F. Carazzolle, Gonçalo A. G. Pereira, Renata M. Baroni

## Abstract

*Moniliophthora perniciosa* and *Moniliophthora roreri* are hemibiotrophic fungi that harbor a large number of Pathogenesis-Related 1 genes, many of which are induced in the biotrophic interaction with *Theobroma cacao.* Here, we provide evidence that the evolution of PR-1 in *Moniliophthora* was adaptive and potentially related to the emergence of the parasitic lifestyle in this genus. Phylogenetic analysis revealed conserved PR-1 genes, shared by many Agaricales saprotrophic species, that have diversified in new PR-1 genes putatively related to pathogenicity in *Moniliophthora*, as well as in recent specialization cases within both species. PR-1 families in *Moniliophthora* with higher evolutionary rates exhibit induced expression in the biotrophic interaction and positive selection clues, supporting the hypothesis that these proteins accumulated adaptive changes in response to host-pathogen arm race. Furthermore, we show that the highly diversified *MpPR-1* genes are not induced by two phytoalexins, suggesting detoxification might not be their main function as proposed before.

## Introduction

Pathogenesis Related-1 (PR-1) proteins are part of CAP (cysteine-rich secretory proteins, antigen 5, and pathogenesis-related 1) superfamily, also known as SCP/TAPS proteins (sperm-coating protein/Tpx-1/Ag5/PR-1/Sc7), and are present throughout the eukaryotic kingdom (Cantacessi et al., 2009; Gibbs et al., 2008). In plants, PR-1 proteins are regarded as markers of induced defense responses against pathogens (van Loon et al., 2006). These proteins have also been ascribed roles in different biological processes in mammals, insects, nematodes and fungi, including reproduction, cellular defense, virulence and evasion of the host immune system (Asojo et al., 2005; Chalmers et al., 2008; Ding et al., 2000; Gao et al., 2001; Hawdon et al., 1999; Lozano-Torres et al., 2014; Prados-Rosales et al., 2012; Schneiter & Di Pietro, 2013; Zhan et al., 2003). In *Saccharomyces cerevisiae*, Pry proteins (*Pathogen related in yeast*) bind and export sterols and fatty acids to the extracellular medium, an activity that has also been demonstrated for other proteins of the CAP superfamily through functional complementation assays (Choudhary & Schneiter, 2012; Darwiche, Mène-Saffrané, et al., 2017; Darwiche & Schneiter, 2016; Gamir et al., 2017; Kelleher et al., 2014).

The basidiomycete fungi *Moniliophthora perniciosa* and *Moniliophthora roreri* are hemibiotrophic phytopathogens that cause, respectively, the Witches’ Broom disease (WBD) and Frosty Pod Rot of cacao (*Theobroma cacao*). Currently, three biotypes are recognized for *M. perniciosa* based on the hosts that each one is able to infect. The C-biotype infects species of *Theobroma* and *Herrania* (Malvaceae); the S-biotype infects plants of the genus *Solanum* (e.g., tomato) and *Capsicum* (pepper); and the L-biotype is associated with species of lianas (Bignoniaceae), without promoting visible disease symptoms (Evans, 1978; Evans, 2007; Purdy & Schmidt, 1996).

With the genome and transcriptome sequencing of the C-biotype, 11 *PR-1*-like genes, named *MpPR-1a* to *k*, were identified in *M. perniciosa* (Teixeira et al., 2012). Interestingly, many of these genes are upregulated during the biotrophic interaction of *M. perniciosa* and *T. cacao*, which constitutes a strong indication of the importance of these proteins in the disease process (Teixeira et al., 2012; Teixeira et al., 2014). In this context, efforts have been made to elucidate the role of these molecules during the interaction of *M. perniciosa* with cacao, such as the determination of the tridimensional structure of MpPR-1i (Baroni et al., 2017) and the functional complementation of *MpPR-1* genes in yeast *Pry* mutants (Darwiche et al., 2017). These studies revealed that seven MpPR-1 proteins display sterol or fatty acid binding and export activity, suggesting that they could function as detoxifying agents against plant lipidic toxins (Darwiche et al., 2017).

Despite these advances, studies with a deeper evolutionary perspective have not yet been performed for MpPR-1 proteins. Evolutionary analysis can be an important tool for the inference of gene function and the identification of mechanisms of evolution of specific traits, such as pathogenicity. Genes that are evolving under negative selection pressures are likely to play a crucial role in basal metabolism (Oleksyk et al., 2010). On the other hand, genes that are evolving under positive selection may have changed to adjust their function to a relatively new environmental pressure (Manel et al., 2016). Thus, it can be hypothesized that *Moniliophthora* PR-1 might have accumulated adaptive substitutions in response to selective pressures related to a pathogenic lifestyle, and the analysis of these substitutions may reveal protein targets and specific codons that are potentially important for the pathogenicity in *Moniliophthora*.

In this study, we performed a two-level evolutionary analysis of *Moniliophthora PR-1* genes: (i) their macroevolution in the order Agaricales, which consists mainly of saprotrophic fungi, being the *Moniliophthora* species one of the few exceptions; (ii) and their microevolution within *M. perniciosa* and its biotypes that differ in host-specificity. By characterizing PR-1 proteins encoded by 22 *Moniliophthora* genomes, reconstructing their phylogenetic history and searching for evidence of positive selection, we identified an increased diversification in these proteins in *Moniliophthora* that is potentially related to its pathogenic lifestyle, as supported by expression data, and also presents cases of species-specific and biotype-specific diversification.

## Results

### Characterization of PR-1 gene families in *Moniliophthora*

Previous work had already reported the identification of 11 *PR-1*-like genes in the genome of *M. perniciosa* isolate CP02 (C-biotype), which were named *MpPR-1a* to *MpPR-1k* (Teixeira et al., 2012). Likewise, 12 *PR-1*-like genes were identified in the genome of *M. roreri* (MCA2977) (Meinhardt et al., 2014). With the sequencing and assembly of 18 additional genomes of *M. perniciosa* isolates and other 4 genomes of *M. roreri* isolates, it was possible to characterize the *PR-1* gene families in the different biotypes of *M. perniciosa* and in its sister species *M. roreri* in order to look for similarities and differences at the species and biotype levels. Figure 1.A shows the number of genes identified as *PR-1* per isolate.

**Figure 1.**
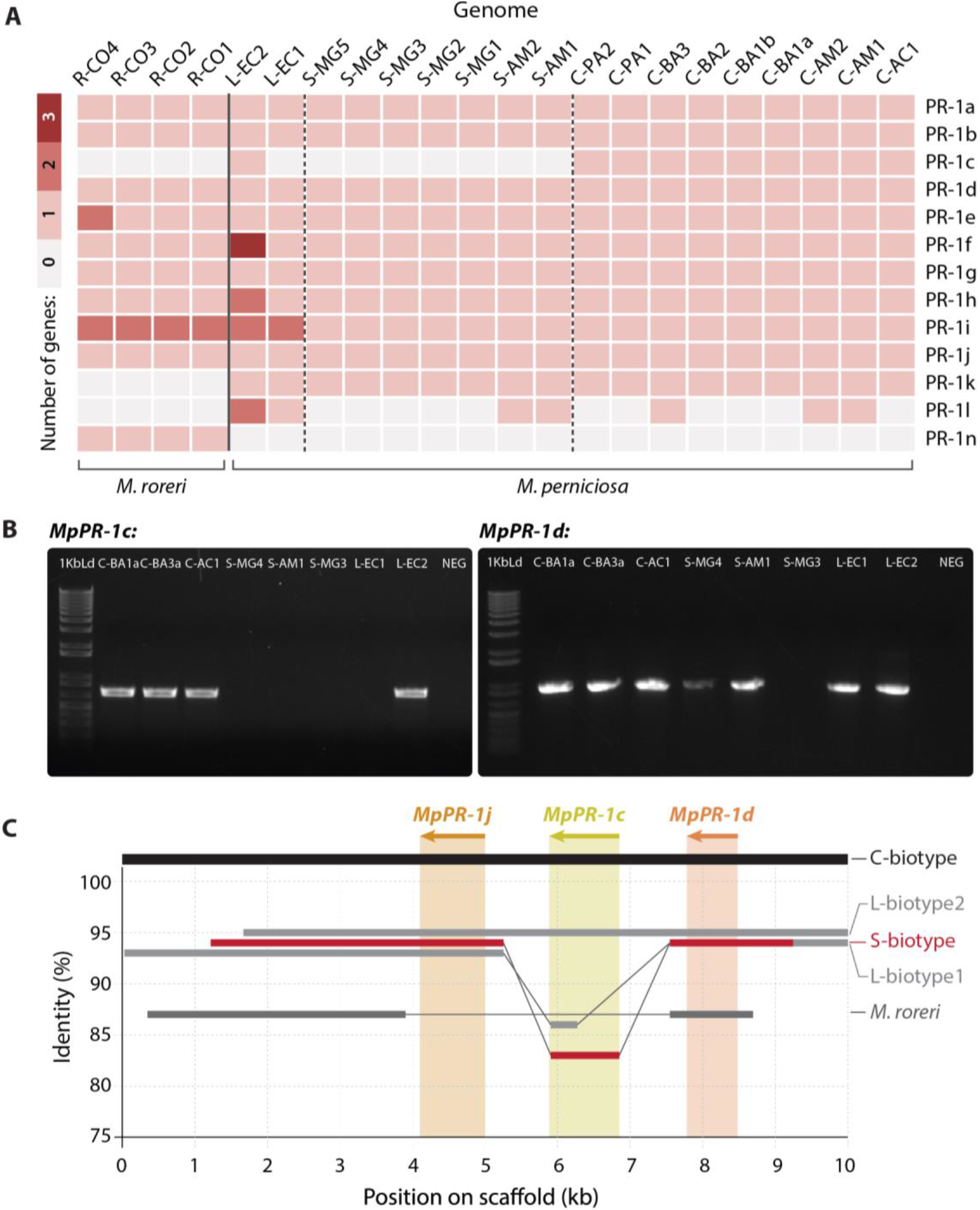
Characterization of PR-1 gene families in *M. perniciosa and M. roreri* genomes. **A.** Heatmap of the number of gene copies per family of *PR-1-like* candidates per *Moniliophthora* isolate. Identification of genomes are in columns and PR-1 family names are in rows. **B.** Amplification by PCR of *MpPR-1c* and *MpPR-1d* genes in the genomic DNA of eight *M. perniciosa* isolates. 1Kb Ld = 1 Kb Plus DNA Ladder (Invitrogen), Neg = PCR negative control (no DNA). Expected fragment sizes were 687 bp for *MpPR-1c* and 902 bp for *MpPR-1d*. **C.** Synteny analysis of a 10 Kb portion of the genome where the *MpPR-1j, c*, *d* genes are found in the three biotypes of *M. perniciosa* and *M. roreri*. The genomes analyzed were C-BA3, S-MG2, R-CO2, L-EC1 and L-EC2. Only identity above 75% to the C-biotype reference is shown.

The examination of orthogroups containing *PR-1-like* hits revealed that the PR-1i orthogroup has the highest number of duplications with two copies in *M. roreri* and in L-biotype. Moreover, a new PR-1-like orthogroup with seven candidates that are more similar to *MpPR-1i* (67% identity) was found. This newly identified gene was named “*MpPR-1l*” and was not found in *M. roreri*. It has the same number and structure of introns and exons as the *MpPR-1i* gene and they are closely located in the same scaffold, which is evidence of a duplication event within *M. perniciosa*. Interestingly, the sequences corresponding to *MpPR-1l* in the five S-biotype isolates from Minas Gerais were found in another orthogroup, in which *MpPR-1l* was fused to the adjacent gene in the genome (a putative endo-polygalacturonase gene containing the IPR011050 domain: Pectin lyase fold) with no start codon found between the two domains. Furthermore, we found that the *MpPR-1i* gene and, consequently, its predicted protein is truncated in almost all S-biotype isolates from MG (except for S-MG2) (Figure 4).

Examining these gene families to look for other putatively species-specific *PR-1* in *Moniliophthora*, we observed that the *MpPR-1k* and *MpPR-1c* genes are not found in the *M. roreri* genomes analyzed in this work, while *MrPR-1n* constitutes an exclusive family in this species. The *MrPR-1o* gene previously identified by Meinhardt et al. (2014) was not predicted in any genome as a gene in this work. The protein sequence of *MrPR-1o* has higher identity with *MrPR-1j* (70%), *MpPR-1j* (66%) and *MpPR-1c* (56%), but it is shorter than all 3 protein sequences and does not have a signal peptide like other PR-1 proteins, which suggested that *MrPR-1o* is a pseudogenized paralog of *MrPR-1j.*

The absence of *MpPR-1c* in all S-biotype genomes suggested that this gene could be biotype-specific within *M. perniciosa*, however, it was predicted in the L-biotype genome L-EC2. Therefore, we sought to confirm the presence or absence of this gene in different *M. perniciosa* by PCR amplification and synteny analysis. The absence of *MpPR-1c* in the S-biotype isolates and in the L-biotype L-EC1 was confirmed, as well as its presence in L-EC2 (Figure 1.B). We also amplified the *MpPR-1d* gene, which was predicted in all genomes, in almost all tested isolates, except for S-MG3 because of a mismatch in the annealing regions of both primers. Even though our PCR results indicated that *MpPR-1c* is not present in the S-biotype, synteny analysis of the genome region where Mp*PR-1j-c-d* are found in tandem (22) revealed that, in fact, *MpPR-1c* is partially present in these S-biotype genomes (Figure 1.C), suggesting again that the duplication event of PR-1j occured in the ancestral of *Moniliophthora* but this paralog was also pseudogenized in the evolution of the S-biotype.

### PR-1 genes evolution along the Agaricales order

To study the macroevolution of PR-1 proteins, we identified orthologous sequences of genes encoding PR-1-like proteins in 16 genomes of species from the Agaricales order, including 3 selected *M. perniciosa* genomes (one of each biotype) and 1 *M. roreri* genome for comparisons. The phylogenetic reconstruction of Agaricales PR-1 proteins revealed a basal separation of two major clades, hereafter called clade1 and clade2 (Figure 2.A).

**Figure 2.**
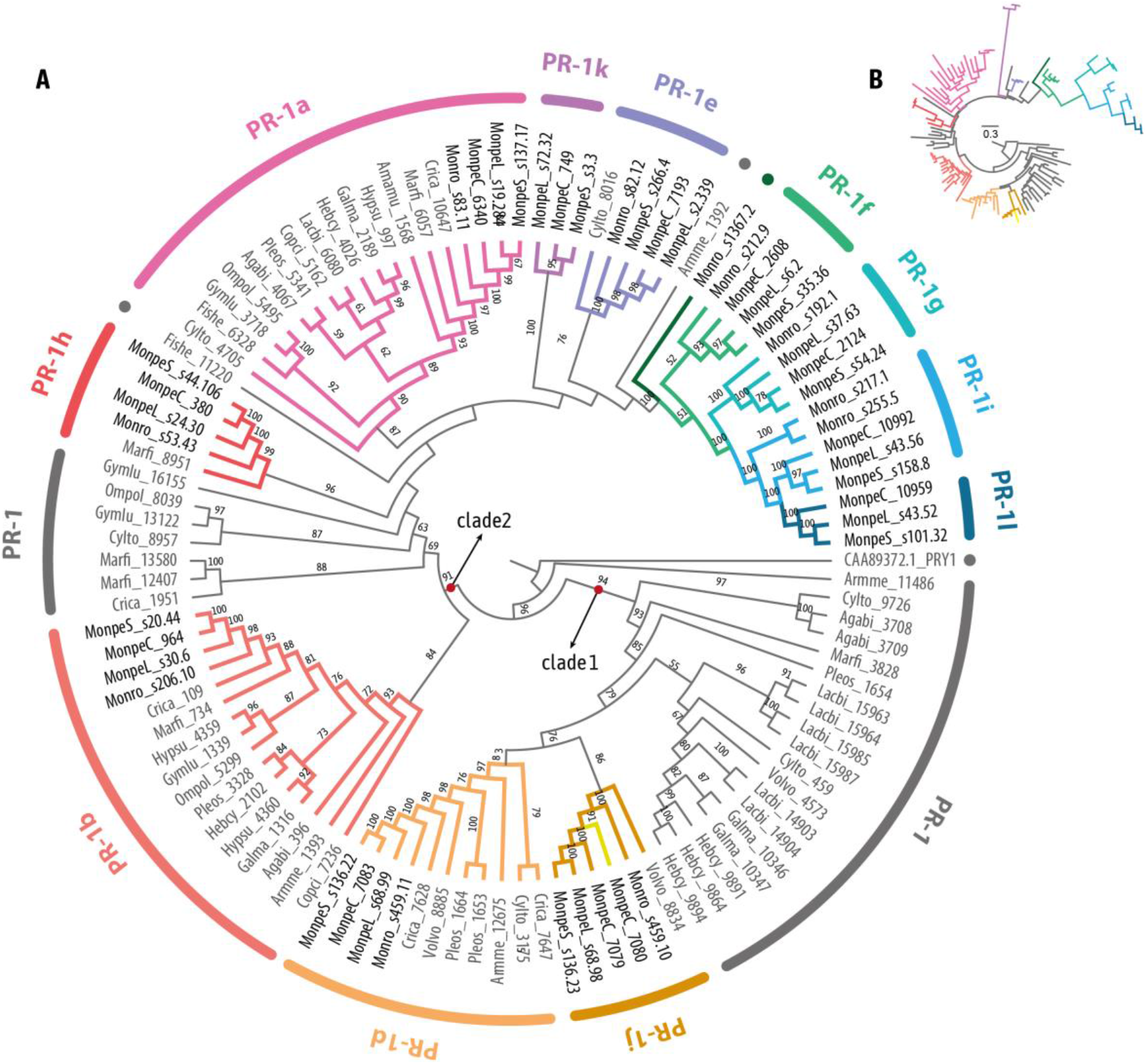
Phylogenetic cladogram of PR-1 proteins in Agaricales (Basidiomycota). **A.** Phylogenetic relationships were inferred by maximum likelihood and branch support was obtained using 1000 bootstraps. Only branch support values greater than 70 are shown. PR-1c is indicated as a yellow branch inside the PR-1j clade, and PR-1n is indicated with a dark green dot and branch. Proteins with ancestral divergence to more than one family were named with the letters of the derived families. Full species names are in SM1. The branch lengths were dimensioned for easy visualization. **B.** The same phylogeny shown in “A” is presented with the respective branch lengths augmented for the *Moniliophthora* specific PR-1 genes.

The first PR-1 clade includes most Agaricales species outside *Moniliophthora* showing *PR-1* genes that diverged early in the phylogeny, before the appearance of PR-1a-l-n orthologues. From this first clade including the early diverged PR-1 proteins, subsequently diverged PR-1d and PR1-j. The separation between PR-1d and j proposed for *Moniliophthora* only occurs in *Volvariella volvacea*, while for all other species, paralogous of PR-1d diverged early. In *Moniliophthora*, PR-1j and PR-1d have more recent common ancestors, and the only paralogous of these *Moniliophthora* PR-1s originates from a possible duplication of *MpPR-1j* in the C-biotype of *M. perniciosa*, which was previously named *MpPR-1c*, therefore exclusive to this species and biotype.

The second clade includes all other PR-1 families and other Agaricales PR-1s that do not have a common ancestor with a single PR-1 from *Moniliophthora.* Clade2 is divided into 2 subclasses in its base, one of them composed of the PR-1b clade, which is distributed among 14 species. In the second subgroup, PR-1a shows a common ancestor in a total of 16 species, being the most common PR-1 here. The great diversification of a PR-1a-like ancestor in *Moniliophthora* resulted in the formation of at least 4 new and exclusive *PR-1* genes (*k*, *g*, *i, l*) with high evolutionary rates reflected on the branch lengths (Figure 2.B). PR-1n showed a putative ortholog in *Armilaria mellea*, the only other plant pathogen in the Agaricales dataset, however, this connection has low branch support.

### Recent diversification of PR-1 genes in *Moniliophthora*

Through the investigation of the phylogenetic history of PR-1 proteins within 22 *Moniliophthora* isolates (Figure 3), we found that the previous classification of *MpPR-1a* to *k* and *MrPR-1n* represents monophyletic clades in the tree, except for *MpPR-1c* which is a recent paralogous of *MpPR-1j*. The evolution of *Moniliophthora* PR-1 also reflected the basal divergence between two large clades as observed in the Agaricales PR-1 tree (Figure 2). The only incongruence between the two phylogenetic trees is the relative position of PR-1k, which appeared after the divergence of PR-1a in Agaricales and before it in *Moniliophthora*. This incongruence may be due to the extreme differentiation of PR-1k, with longer branch lengths in *Moniliophthora*, being its position on the Agaricales tree more reliable.

**Figure 3.**
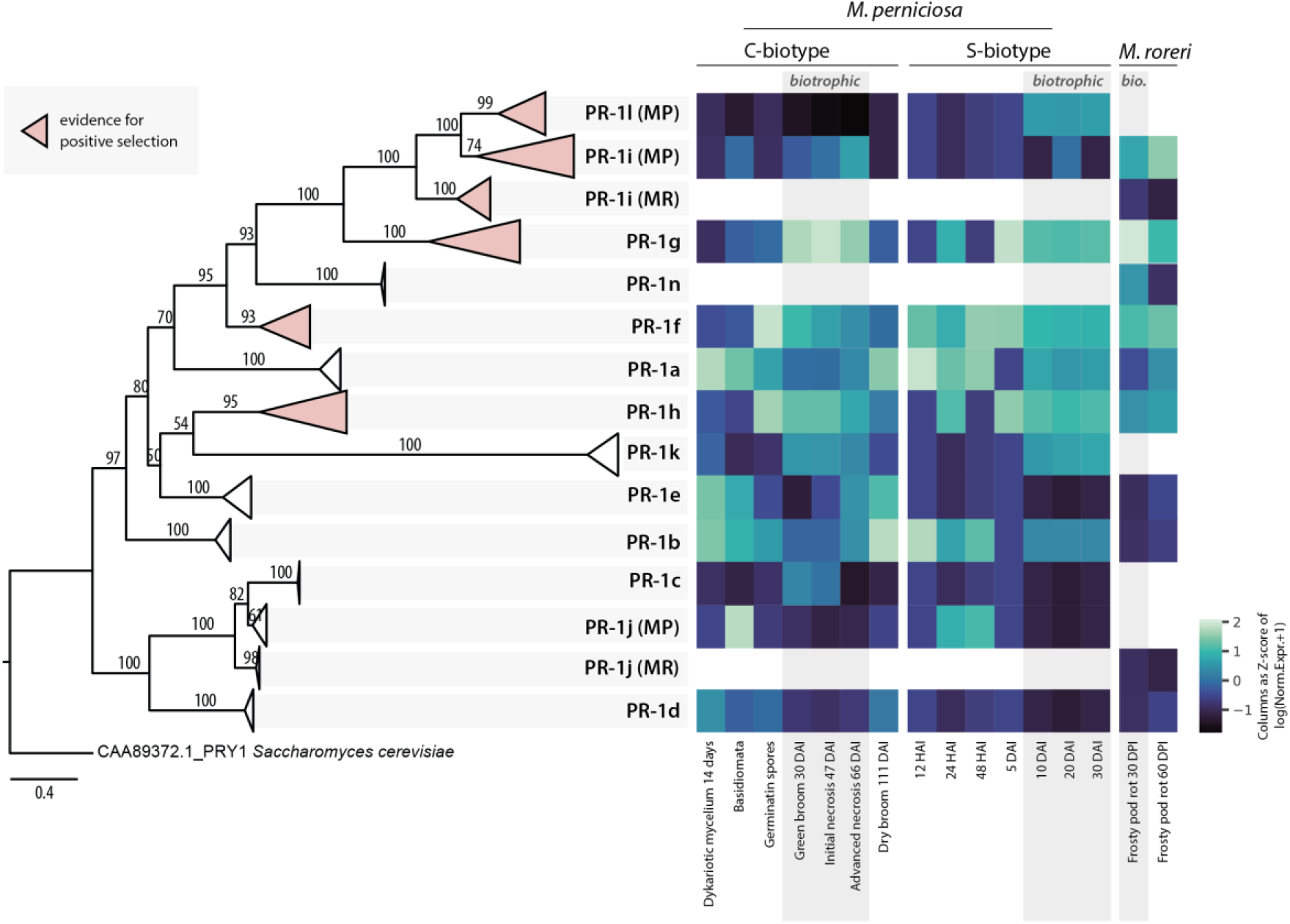
Phylogenetic reconstruction of PR-1 proteins in *Moniliophthora* and heatmap of expression Z-scores of PR-1. Phylogenetic relationships of PR-1 proteins from 18 *M. perniciosa* and 4 M. *roreri* isolates were inferred by maximum likelihood and branch support was obtained using 1000 bootstraps. The PRY1 protein of *Saccharomyces cerevisiae* was used as an outgroup. Clades filled with pink color represent PR-1 families with evidence of positive selection. A version of this tree with non-collapsed branches can be found in Supplementary Figure 1. For each PR-1 family, the Z-score of log transformed expression levels of *MpPR-1* and *MrPR-1* from transcriptomic data was calculated for conditions (columns) and plotted as a heatmap. The heatmap includes *MpPR-1* data from seven conditions of the C-biotype of *M. perniciosa* from the Witches’ Broom Transcriptomic Atlas, 7 different time points of S-biotype infection in MicroTom tomato plants, and two conditions of *M. roreri* infection in cacao pods (frosty pod rot). Conditions highlighted with a grey background indicate the biotrophic stage of the plant-pathogen interaction.

Among the PR-1 families, the proteins with the greatest number of changes in the tree are PR-1g, i, and k, which are exclusive PR-1s in the genus *Moniliophthora*, as pointed out by the previous phylogenetic analysis. In addition, PR-1h also showed a greater branch length than the others, being a family of PR-1s only shared between *Moniliophthora* and *Marasmius* in the Agaricales PR-1 tree (Figure 2). PR-1c also presented a large number of changes in relation to its ancestor PR-1j. This greater number of changes in these MpPR-1s, and their exclusive presence in comparison to the other Agaricales, indicate a recent potential adaptive process of diversification of these proteins in *Moniliophthora*.

### Positive selection shaping PR-1 families in *Moniliophthora*

Based on the observations of high diversification of PR-1 families within *Moniliophthora*, we hypothesized that positive selection could be shaping these proteins either in the C-biotype or in the S-biotype. To test this hypothesis, we tested the branch-sites evolutionary model for each PR-1 family. None of these tests brought evidence of positive selection in any PR-1 family for the C-biotype branches. For the S-biotype branches, a signal of positive selection was detected for PR-1g on one site of the protein sequence.

Considering that the existence of *M. perniciosa* biotypes are very recent in the evolutionary timescale and that C-biotype itself has almost no genetic variation among its sequences, which makes it very difficult to apply separate dN/dS tests, we tested both C- and S- biotypes together. We tested the hypothesis that there was a single selective pressure shaping PR-1 families throughout the *M. perniciosa* and *M. roreri* evolution regardless of the biotype, using the site model test. In these tests, sites with positive selection signs were detected in five families (PR-1f, g, h, i, l) (Table 2). The PR-1n family was not included in these tests because all sequences were identical.

**Table 2.** Omega (dN/dS) values and protein sites (amino acid: position) detected with significant probability of positive selection for each PR-1 family in *Moniliophthora*.

| Family | Omega | Sites under positive selection (p>0.95) |
| --- | --- | --- |
| PR-1a | 4.12 | None |
| PR-1b | 2.31 | None |
| PR-1c | 1 | None |
| PR-1d | 2.07 | None |
| PR-1e | 1 | None |
| PR-1f | 2.74 | K: 49, S: 157 |
| PR-1g | 7.05 | P: 211, P: 234, A: 242, S: 260, S: 271 |
| PR-1h | 4.33 | S: 78, Y: 107, P: 141, S: 155, E: 187, D: 206, L: 209, M: 224, R: 259, Q: 267 |
| PR-1i | 4.38 | T: 40, Q: 54, D: 56, R: 118, K: 132, A: 141, L: 150 |
| PR-1l | 6.39 | Q: 65, Q: 76, -: 164, N: 176, R: 192, K: 193, E: 194, F: 196 |
| PR-1j | 2.76 | None |
| PR-1k | 2.50 | None |

PR-1g stands out for having the highest omega and for being one of the most expressed genes during the green broom phase (Teixeira et al., 2014). Three of the codons under positive selection are part of the ‘keke’ domain, which is possibly involved in the interaction with divalent ions or proteins (Teixeira et al., 2012). Among the sites detected under positive selection for PR-1i, one is found in the caveolin binding motif (CBM), an important region for binding to sterols, and another site is in the alpha-helix 1, which together with alpha-helix 4 form the cavity for ligation to palmitate (Baroni et al., 2017) (Figure 4).

**Figure 4.**
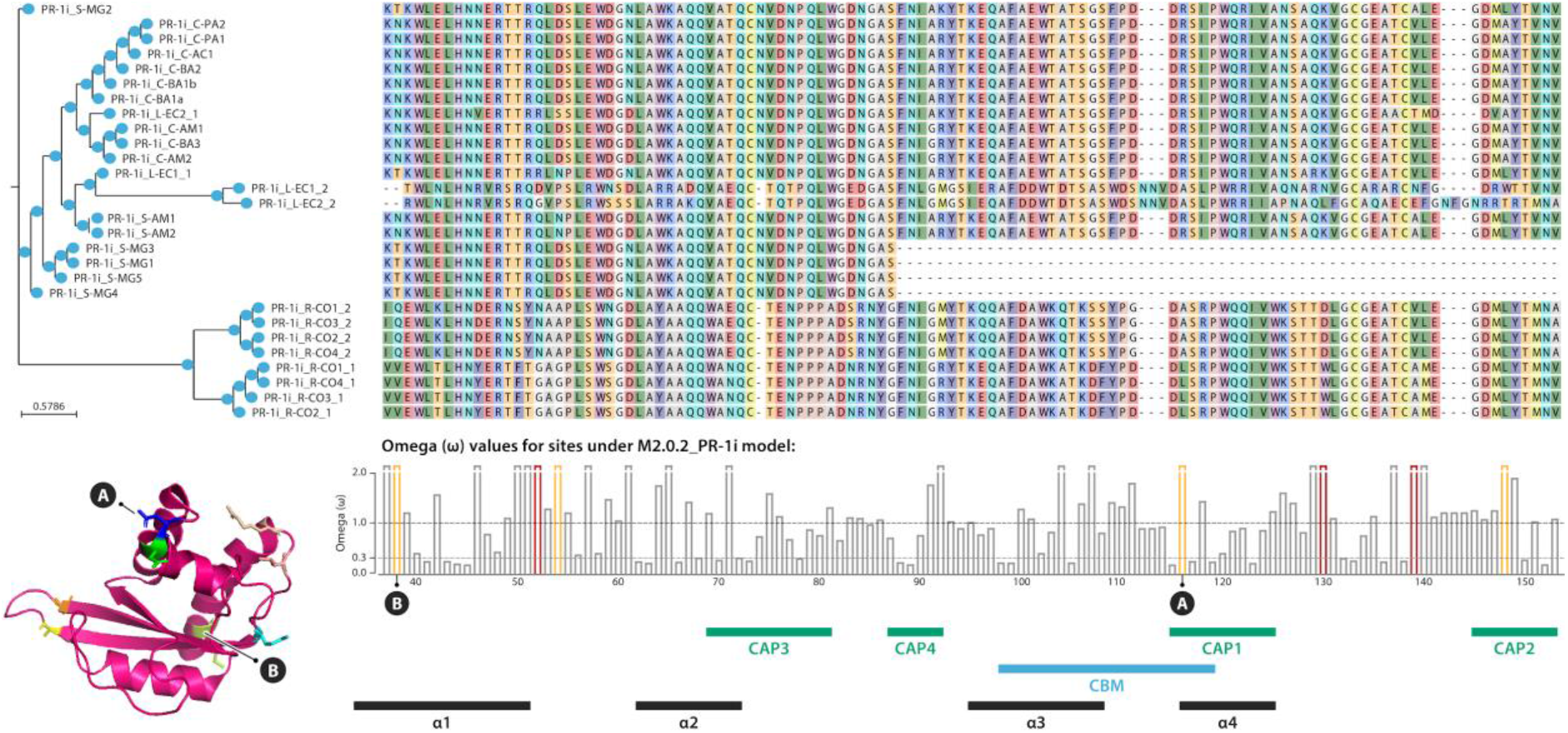
Sequence alignment and phylogeny of PR-1i proteins in *Moniliophthora* isolates. Only a slice of the middle portion of the alignment is shown to highlight the sites with positive selection signs, indicated by red (p-value ≤ 0.01) or orange (p-value ≤ 0.05) bars in the bar chart of omega values below the alignment. Below de bar chart, annotations indicate the locations along the sequence of the CAP domains, caveolin-binding motif (CBM) and alpha-helices (α). On the 3D crystal structure of MpPR-1i protein (PBD:5V50) and on the bar chart, “A” indicates the site under positive selection detected in the CBM and “B” indicates the site under positive selection in alpha-helix 1.

Although the two exclusive PR-1 families of *M. perniciosa*, PR-1c and PR-1k, do not present evidence of positive selection, both revealed processes of diversification in the PR-1 phylogenies. It is possible that these families have also undergone selective pressures in their evolution, but the short time of evolution of *M. perniciosa* in relation to the genus has reduced the accuracy of the dN/dS tests in these exclusive families.

### Adaptive evolution of PR-1 is reflected on expression data

It has already been shown that *MpPR-1* genes of the C-biotype have distinct expression profiles in several different conditions of the WBD Transcriptome Atlas, which were also confirmed by quantitative RT-PCR (Teixeira et al., 2012; Teixeira et al., 2014). *MpPR-1a*, *b*, *d*, *e* are ubiquitously expressed during the necrotrophic mycelial stage, while *MpPR-1j* is mainly expressed in primordia and basidiomata. Six *MpPR-1*s are highly and almost exclusively expressed during the biotrophic stage of WBD: *MpPR-1c*, *f, g*, *h*, *i*, *k* (Teixeira et al., 2012). However, in contrast to *MpPR-1i*, the newly discovered *MpPR-1l* is not expressed in any of the conditions analyzed, suggesting that this gene may not be functional in the C-biotype.

Expression data of *MpPR-1* genes from the S-biotype during a time course of the biotrophic interaction with MT tomato revealed that *MpPR-1f*, *g*, *h*, *l*, and *k* are highly expressed during 10-30 days after infection (d.a.i.) (Figure 3). These results are similar to the expression profile verified for the C-biotype during the biotrophic interaction with *T. cacao*, with the exceptions that *MpPR-1c* is absent in the S-biotype and, instead of *MpPR-1i*, *MpPR-1l* is expressed during tomato infection. S-biotype *MpPR-1*s are highly expressed starting at 10 d.a.i., which is usually when the first symptoms of stem swelling are visible in MT tomato (Deganello et al., 2014). *MpPR-1a* and *b* appear to have ubiquitous expression profiles since they show similar expression levels in almost all conditions. *MpPR-1j, d*, *e*, and *i* did not show significant expression in these libraries. Because *MpPR-1i* is truncated in the S-MG1 genome, quantification of expression was also done with S-MG2 as a reference, since it has a complete *MpPR-1i* gene. However, we still obtained the same expression profiles as S-MG1 for all *MpPR-1*. This could suggest that *MpPR-1l* is expressed in S-biotype even with a fusion to the adjacent gene. In the C-biotype, this adjacent gene is only expressed during the biotrophic interaction.

In *M. roreri*, it has been previously reported that *MrPR-1n*, *MrPR-1g* and *MrPR-1i2* were upregulated in samples from the biotrophic phase (30 days post infection of pods), *MrPR-1d* was upregulated in the necrotrophic phase (60 days post infection of pods) and five other *MrPR-1* were constitutively expressed under these conditions (Meinhardt et al., 2014). The heatmap in Figure 3 shows that similar to *M. perniciosa’s PR-1* expression profile, *MrPR-1g* is the most expressed *PR-1* gene during the biotrophic stage. Moreover, while *MrPR-1h* and *MrPR-1f* were not differentially expressed when comparing the biotrophic and necrotrophic stages, they also showed higher expression when compared to other *MrPR-1*s that belong to the conserved families.

### Expression of recently diversified MpPR-1 was not induced by plant antifungal compounds

It has been previously demonstrated that CAP proteins of *M. perniciosa* bind to a variety of small hydrophobic ligands with different specificities. Thus, it has been suggested that the *MpPR-1* genes induced in the biotrophic interaction could function in the detoxification of hydrophobic molecules produced by the host as a defense strategy (Darwiche et al., 2017). In this context, we investigated if *MpPR-1* genes, especially the ones induced in WBD (*c, f, h, i, k, g*), are differentially expressed by the presence of the plant antifungal compounds eugenol or α-tomatin, which are similar to sterol and fatty acids, respectively. However, when the necrotrophic mycelia of *M. perniciosa* was treated with eugenol, only *MpPR-1e, k, d* were up-regulated, while *MpPR-1f* was down-regulated in α-tomatin-treated samples (Figure 5). In all samples, among all *MpPR-1* genes, *MpPR-1a* and *MpPR-1b* had the highest expression levels, while *MpPR-1c, i, g, j* have the lowest (TPM ≤ 2).

**Figure 5.**
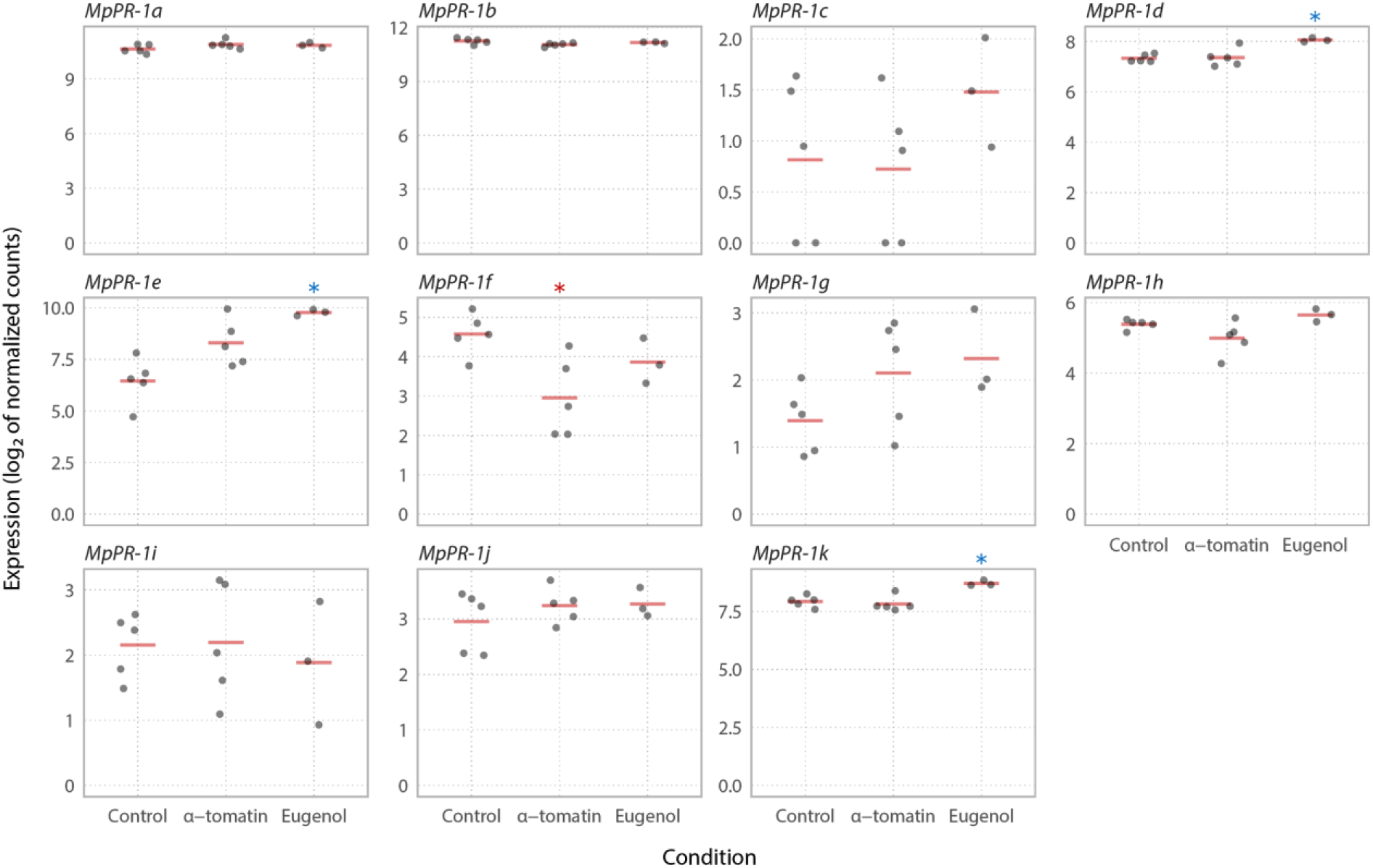
Expression profile of *MpPR-1* genes in response to two plant antimicrobial compounds. The necrotrophic mycelium of *M. perniciosa* C-biotype (C-BA1a) was grown in liquid media in the presence of eugenol, α-tomatin or DMSO (mock condition) for 7 days. The expression values (Log2 transformed) for each *MpPR-1* were obtained by RNA-Seq and subsequent quantification of read counts and between-sample normalization using size factors. Red bars indicate the mean of expression values within a group of replicates. Asterisks indicate that *MpPR-1e, k, d, f* are differentially expressed (s-value<0.005) when compared to the mock condition, with blue asterisk indicating up-regulation and red indicating down-regulation.

## Discussion

### The evolution of PR-1 and the emergence of pathogenicity among saprotrophs

The plant pathogen *M. perniciosa* has at least 11 genes encoding *PR-1*-like secreted proteins, which were previously identified and characterized in the genome of the C-biotype CP02 isolate (Teixeira et al., 2012). Many of these genes were shown to be highly expressed during the biotrophic interaction of *M. perniciosa* and cacao, suggesting that MpPR-1 proteins have important roles during this stage of WBD. *M. perniciosa* has two other known biotypes (S and L) that differ in host specificity and virulence, the closest related species *M. roreri* that also is a *T. cacao* pathogen, and other nine *Moniliophthora* species: one described as a non-pathogenic grass endophyte (Aime & Phillips-Mora, 2005), three of biotrophic/parasitic habit (Niveiro et al., 2020), and five species of unascertained lifestyle, found in dead or decaying vegetal substrates (Kerekes et al., 2009; Kropp & Albee-Scott, 2012; Takahashi, 2002). Because the majority of *Moniliophthora* related fungi in the Agaricales order are saprotrophs, the occurrence of parasitic *Moniliophthora* species raises the question about the emergence of biotrophic/parasitic lifestyle in this lineage of Marasmiaceae (Niveiro et al., 2020; Teixeira et al., 2015). The evolutionary scenario of host-pathogen arms race that emerges through the diversification of the *Moniliophthora* genus in the Agaricales order and of host-specific biotypes in *M. perniciosa* isolates, is especially suitable for the study of adaptive evolution in pathogenicity-related genes. Besides, the knowledge on putative adaptations gained through pathogen evolution are also specially interesting for further development of strategies against the pathogen. Based on our findings, Figure 6 presents a model for the adaptive evolution of PR-1 proteins in *Moniliophthora*.

**Figure 6.**
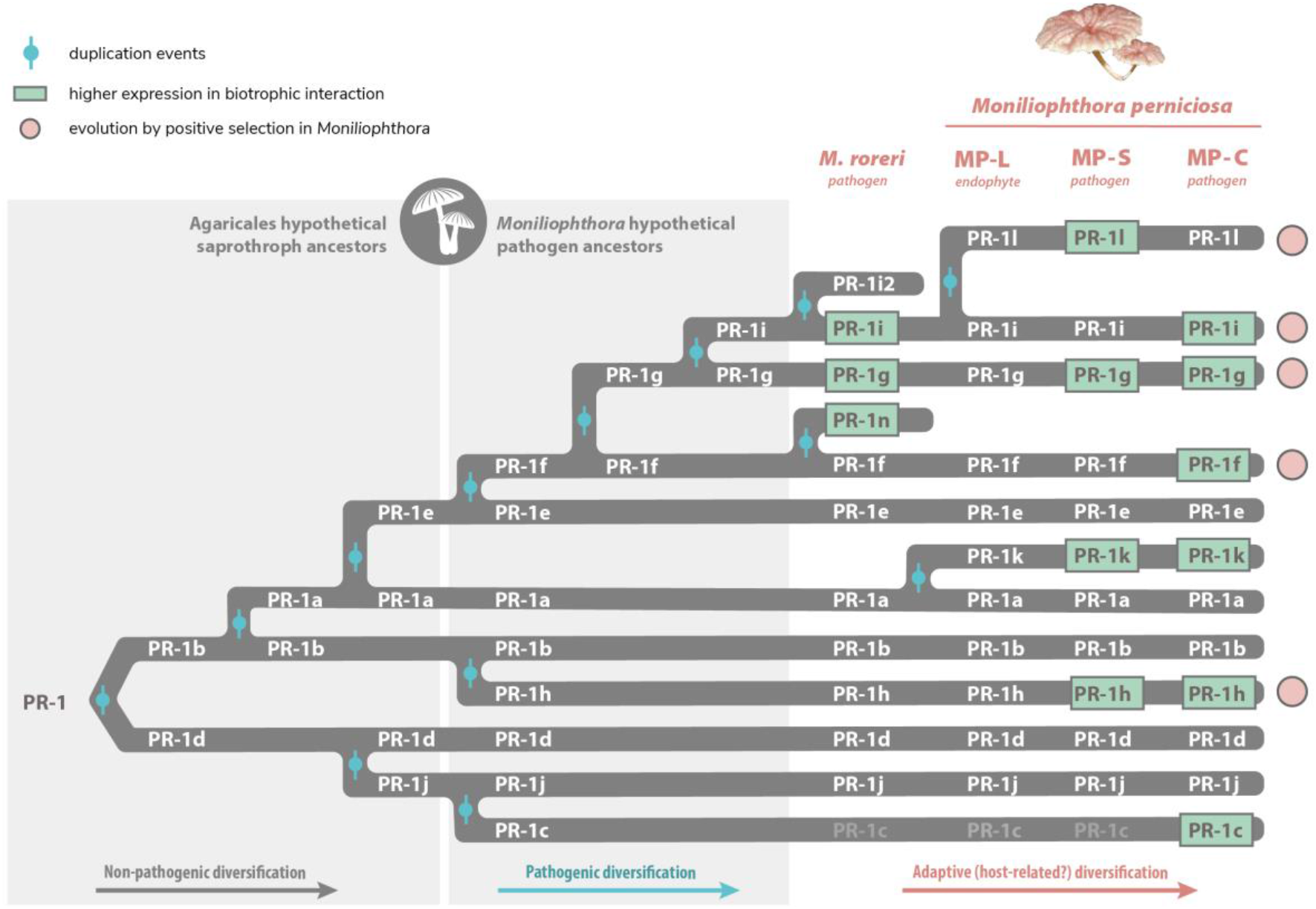
Proposed model for the adaptive evolution of PR-1 proteins in *Moniliophthora* towards the pathogenic lifestyle. All *Moniliophthora* PR-1 proteins derived independently from two ancient clades (PR-1b-like and PR-1d-like) within the Agaricales order, as indicated in PR-1 phylogeny. The subsequent diversification of PR-1a and PR-1e from PR-1b, and PR-1j from PR-1d, occured in putative saprotroph lineages before the divergence of *Moniliophthora* genus, suggesting a diversification not related to pathogenicity. Within *Moniliophthora* hypothetical pathogenic ancestors, five other PR-1 proteins were derived (c from j, h from b, f-g-i from e) and most of these new lineages showed evidence of positive selection in *M. perniciosa* samples (indicated by pink circles). New PR-1 copies (n and i2 in *M. roreri*, l and k in *M. perniciosa*) diverged within *M.* species. Recently diversified PR-1 genes in *Moniliophthora,* not only show an elevated rate of evolution and positive selection evidence but are also predominantly expressed during the biotrophic interaction (indicated by green highlights). This supports the hypothesis that these proteins accumulated adaptive changes related to pathogen lifestyle that might also contribute to the host specialization observed in *Moniliophthora* species and biotypes.

Through the characterization of the evolution of PR-1 proteins in Agaricales, we observe that at least one copy of PR-1 is present in all the sampled fungi, with most of the Agaricales species encoding between 1 and 7 PR-1 proteins. This is in contrast with *Moniliophthora* species, which encode among 10-12 proteins. *Moniliophthora* PR-1 proteins are derived independently from both ancient clades in the Agaricales gene tree. The longer branch-lengths in PR-1 families exclusive to *Moniliophthora* along with the evidence of evolution under positive selection identified in independently diverged clades suggest that the diversification of PR-1 in the genus was adaptive and related to its pathogenic lifestyle. The accentuated adaptive evolution of PR-1 in *Moniliophthora* is not only reflected in the genomic evolution of these genes, but also in their expression context. *PR-1c*, *f*, *g*, *h*, *i*, *k*, *l*, *n* are upregulated during the biotrophic interaction, while *PR-1*s that are also conserved in other Agaricales species are mainly expressed in mycelial stages of *M. perniciosa* (*MpPR-1a*, *b*, *d*, *e)*. Most Agaricales species are not plant pathogens, which is also an evidence that PR-1 in *Moniliophthora* diverged from a few ancestral PR-1s that are related to basal metabolism in fungi and have been evolving under positive selective pressure possibly because of a benefit for the biotrophic/pathogenic lifestyle.

The emergence of SCP/TAPs proteins as pathogenicity factors has been reported in other organisms, such as the yeast *Candida albicans* and the filamentous fungus *Fusarium oxysporum* (Braun et al., 2000; Prados-Rosales et al., 2012). Even though their specific function and mode of action may be different and remains to be characterized in plant pathogens, the recent evolution of these proteins towards their pathogenic role in the *Moniliophthora* genus could have contributed for the transition from a saprotrophic to parasitic lifestyle. Accelerated adaptive evolution evidenced by positive selection signs has also been observed in other virulence-associated genes of pathogenic fungi, such as the genes *PabaA*, *fos-1*, *pes1*, and *pksP* of *Aspergillus fumigatus*, which are involved in nutrient acquisition and oxidative stress response (Fedorova et al., 2008), and several gene families in *C. albicans*, including cell surface protein genes enriched in the most pathogenic *Candida* species (Butler et al., 2009).

### Adaptive evolution of PR-1 within *Moniliophthora* species and its biotypes

The high diversification of PR-1 families observed within *Moniliophthora* was reinforced by our findings of positive selection in families that have also augmented expressions during infection: PR-1f, g, h, i, l. Among these five families, PR-1g and PR-1i are two of the most diversified families in *Moniliophthora* and have a more recent common ancestor with PR-1f than with the other PR-1s, placing this monophyletic clade of PR-1f, g, i as the key one to diversification and adaptive evolution of these proteins in the genus. Many sites that potentially evolved under selective pressure were also found in PR1-h, indicating that a parallel process of adaptive evolution occurred in this family.

Within the diversification of PR-1 in *Moniliophthora*, four cases of putative species-specific evolution of PR-1 families were found: PR-1c, PR-1k and PR-1l in *M. perniciosa* and PR-1n in *M. roreri*. Although vestigial sequences indicate that PR-1c probably emerged as a paralog of PR-1j in the ancestral of both species, it was only kept in the evolution of C-biotype, in which a change in expression profile occurred, thus placing *MpPR-1c* as a case of PR-1 diversification within *M. perniciosa* biotypes and a potential candidate for host specificity. Another candidate for biotype-specific diversification is PR-1l, which diverged from a duplication of PR-1i. Even though PR-1l was found in all three *M. perniciosa* biotypes, it was expressed only in the S-biotype during the biotrophic interaction, instead of PR-1i, which is expressed in *M. roreri* and in the C-biotype. This suggests that the divergence of PR-1i can be host-specific, but further experiments are necessary to clarify if they are either a cause or consequence of *M. perniciosa* pathogenicity.

Even though almost all PR-1 families are present in the genomes of L-biotype isolates, this biotype has an endophytic lifestyle and does not cause visible disease symptoms in their hosts (H. C. Evans, 1978; Griffith & Hedger, 1994). There is no available expression data for the L-biotype, so it is unknown whether their PR-1 genes could have any role related to their lifestyle or these genes are not pseudogenized yet due to a small evolutionary time. Evans (1978) reported that the L-biotype can induce weak symptoms in seedlings of the Catongo variety of *T. cacao*. Therefore, it could be possible that host susceptibility is an important factor for the manifestation of WBD symptoms.

### From basal metabolism to key roles in disease: How PR-1 proteins could have functionally adapted for pathogenicity?

It was previously shown that Pry proteins detoxify and protect yeast cells against eugenol (Darwiche, Mène-Saffrané, et al., 2017) and that MpPR-1 proteins can bind to hydrophobic compounds secreted by plants, indicating that they could antagonize the host defense response (Darwiche, El Atab, et al., 2017). When the necrotrophic mycelia of *M. perniciosa* was cultivated with eugenol, expression of *MpPR-1d* and *MpPR-1k* was up-regulated, which is in agreement with the ability of these two proteins to bind to plant and fungal sterol compounds (Darwiche, El Atab, et al., 2017). *MpPR-1e* expression was also highly induced by eugenol, even though it was previously shown to bind only to fatty acids, but not sterols. Additionally, *MpPR-1f*, which is up-regulated during the biotrophic interaction like *MpPR-1k*, was down-regulated by α-tomatin. Given the above, it appears that *M. perniciosa* does not rely on MpPR-1 for cellular detoxification or this function is not transcriptionally regulated by plant hydrophobic compounds in the necrotrophic mycelia, or even, they could have different roles other than detoxification.

Considering that some PR-1 proteins are not associated with infection and are conserved in other saprotrophic fungi, here we hypothesize that the primary function of PR-1 in fungi can be related to the export of sterols from basal metabolism, such as ergosterol, the most abundant sterol in fungal cell membrane (Mohd et al., 2011; Zhao et al., 2005). PRY of *S. cerevisiae* transports acetylated ergosterol to the plasma membrane (Choudhary & Schneiter, 2012) and MpPR-1d, which belongs to a relatively conserved PR-1 family in Agaricales, can also efficiently bind to ergosterol (Darwiche et al., 2017). Furthermore, ergosterol acts as a PAMP molecule (pathogen-associated molecular pattern) in plants (Nürnberger et al., 2004), resulting in the activation of defense-related secondary metabolites and genes, including plant PR-1s (Kasparovsky et al., 2003; Klemptner et al., 2014; Lochman & Mikes, 2006; van Loon et al., 2006), which are likely to have a role in sequestering sterols from the membranes of microbes (Gamir et al., 2017) and stress signaling (Chen et al., 2014; Chien et al., 2015). Additionally, PR-1 receptor-like kinases (PR-1-RLK) from *T. cacao* are also upregulated on WBD and could be binding to the same ligand of PR-1 (Teixeira et al., 2013). Given that, it is possible that MpPR-1 could have evolved different adaptive roles through neofunctionalization. Besides export of hydrophobic compounds of basal metabolism, they could be acting in the protection of the cell membrane against the disruption caused by antifungal compounds, in the detoxification of hydrophobic compounds like phytoalexins secreted by the host, or it could even be possible that those *MpPR-1s* expressed during infection are sequestering the membrane sterols of the fungus itself in order to prevent detection by a possible ergosterol recognition complex from the host (Khoza et al., 2019), thus compromising the elicitation of plant immunity in a similar fashion of MpChi, a chitinase-like effector that is highly expressed by *M. perniciosa* during the biotrophic stage of WDB (Fiorin et al., 2018).

It has been shown that the ability of MpPR-1 proteins to bind to sterols can be altered by a point mutation in the caveolin binding motif (Darwiche et al., 2017), highlighting the significance of understanding those sites under positive selection that are detected in important regions of the proteins, such as the candidate sites found in the CBM and alpha-helix 1 of PR-1i. These findings are central to learn how changes in the nucleotide or protein sequences could impact binding affinity and function. Even though this is speculative, as the specific role of PR-1 remains unknown, these results can guide further validation experiments and maybe demonstrate another case of adaptive evolution of fungal effectors.

## Conclusions

Based on genomic and transcriptomic data, we presented evidence of adaptive evolution of PR-1 proteins in processes underlying the pathogenic lifestyle in *Moniliophthora*. These results reinforce the power of evolutionary analysis to reveal key proteins in the genomes of pathogenic fungi and contribute to the understanding of the evolution of pathogenesis. Our results indicate a set of PR-1 families that are putatively related to pathogenicity in the genus (PR-1f, g, h, i) and specialization within *M. perniciosa* biotypes (PR-1c, k and l) and *M. roreri* (PR-1n). The positive selection analysis also indicates protein sites that are putatively related to those adaptations. *PR-1* genes and sites with evidence of adaptations are strong candidates for further study and should be evaluated in order to understand how changes in these sites can affect structure, binding affinity and function of these proteins.

## Material and Methods

### Identification of PR-1-like gene families

In this study, we used a dataset of families of genes predicted in 22 genomes of *Moniliophthora* (Filho Tokimatu, 2018) and 16 genomes of other fungal species of the order Agaricales, which were obtained from the Joint Genome Institute (JGI) Mycocosm database (Grigoriev et al., 2014). The *Moniliophthora* genomes included are 7 isolates of the S-biotype (collected at the states of Amazonas and Minas Gerais, in Brazil), 9 isolates of the C-biotype (collected at the states of Amazonas, Pará, and Acre, in Brazil), 2 isolates of the L-biotype (from Colombia) and 4 samples of *M. roreri* (from Colombia). Supplementary File 1 contains the list of species and isolates, their genome identification and source (collection location or reference publication).

To identify candidate *PR-1* gene families, we performed a search for genes encoding the CAP/SCP/PR1-like domain (CDD: cd05381, Pfam PF00188) using the HMMER software (Eddy, 2011). The assignment of protein sequences to families of homologues (orthogroups) was done using Orthofinder (v. 1.1.2) (Emms & Kelly, 2015). In addition, we searched all gene families for families containing *PR-1* candidate genes with Blastp (Camacho et al., 2009) using the known 11 MpPR-1 sequences (Teixeira et al., 2012) as baits, in order to search for possible candidates that were not previously identified and/or that have been wrongly assigned to other orthogroups due to incorrect gene prediction. To verify the presence of the SCP PR1-like/CAP domain (InterPro entry IPR014044) in the sequence, the InterProscan platform (Hunter et al., 2009) was used. All PR-1 candidate sequences identified in this study are deposited in GenBank under accession numbers MW659198 - MW659445.

### Sequence alignment and phylogenetic reconstruction

For the inference of the phylogenetic history of the gene, the protein sequences of the PR-1 homologue families identified in the 22 *Moniliophthora* isolates were aligned with the PRY1 sequence of *S. cerevisiae* (GenBank ID CAA89372.1), which was used as outgroup. Multiple sequence alignments were performed with Mafft (v. 7.407) (Katoh & Standley, 2013) using the iterative refinement method that incorporates local alignment information in pairs (L-INS-i), with 1000 iterations performed. Then, the alignments were used for phylogenetic reconstruction using the maximum likelihood method with IQ-Tree (v. 1.6.6) (Nguyen et al., 2015), which performs the selection of the best replacement model automatically, with 1000 bootstraps for branch support. Bootstraps were recalculated with BOOSTER (v. 0.1.2) for better support of branches in large phylogenies (Lemoine et al., 2018). Likewise, the phylogenetic inference for PR-1 of the Agaricales group of species was performed with the alignment of the homologous proteins identified in the 16 species obtained from Mycocosm, 3 isolates of *M. perniciosa* (C-BA3, S-MG3, L-EC1, one representing each biotype), an isolate of *M. roreri* (R-CO1), and PRY1 of *S. cerevisiae* as the outgroup. To improve alignment quality, trimAl package (Capella-Gutiérrez et al., 2009) was used. For dN/dS analysis, considering each gene family independently, the phylogenetic reconstruction was performed using IQ-Tree (v. 1.6.6) with the multiple local alignment of the protein sequences obtained with Mafft (v. 7.407), and the codon-based alignment of the nucleotide sequences was performed with Macse (v. 2.01) (Ranwez et al., 2018).

### Detection of positive selection signals

To search for genes and regions that are potentially under positive selection in each of the PR-1 families of the 22 isolates, the CodeML program of the PAML 4.7 package (Yang, 2007) was used with the ETEToolkit tool (Huerta-Cepas et al., 2016). CodeML implements a modification of the model proposed by (Goldman & Yang, 1994) to calculate the omega (rate of non-synonymous mutations (dN)/rate of synonymous mutations (dS)) of a coding gene from the multiple alignment sequences and phylogenetic relationships that have been previously inferred.

In order to detect positive selection signals in isolates or specific positions in the sequences, we performed tests with the “branch-site” model, which compares a null model (bsA1) in which the branch under consideration is evolving without restrictions (dN/dS = 1) against a model in which the same branch has sites evolving under positive selection (bsA) (dN/dS > 1) (Zhang et al., 2005). In these tests, those branches that were tested for significantly different evolving rates from the others (foreground branches ωfrg) are marked in the phylogenetic trees – in this case, the branches corresponding to the isolates of the C-biotype or S-biotype. To detect signs of positive selection at specific sites throughout the sequences, regardless of the isolate, we used the “sites” model (M2 and M1, NSsites 0 1 2) to test all branches of the phylogenetic trees.

In both tests, the models are executed several times with different initial omegas (0.2, 0.7, 1.2), and the models with the highest probability are selected for the hypothesis test, in which a comparison between the alternative model and the null model is made through a likelihood ratio test. If the alternative model is the most likely one (p-value <0.05), then the possibility of positive selection (ω>1) can be accepted, and sites with evidence of selection (probability> 0.95) are reported by Bayes Empirical Bayes analysis (BEB) (Zhang et al., 2005).

### Gene amplification and synteny analysis of PR-1c

In order to confirm the presence or absence of *MpPR-1c* and *MpPR-1d* genes in the genomes of *M. perniciosa* isolates, these genes were amplified by polymerase chain reaction (PCR) from isolates C-AC1, C-BA1a, C-BA3, S-AM1, S-MG3, S-MG4, L-EC1 and L-EC2. The necrotrophic mycelia of these isolates was cultivated in 1.7% MYEA media (15 g L^−1^ agar; 5 g L^−1^ yeast extract, 17 g L^−1^ malt extract) at 28°C for 14 days, then harvested and ground in liquid nitrogen for total DNA isolation with the phenol-chloroform method (Sambrook & Russell, 2006). PCRs were performed with primers designed for *MpPR-1c* (F: 5’-GGATCCCGACTTGACAACTCCATCTCG-3’, R: 5’-GAGCTCTCACTCAAACTCCCCGTCATAAT-3’) and *MpPR-1d* (F: 5’-GGATCCCCCTCGCAATGGGTTTTC-3’, R: 5’-GTCGACTCAGTCAAGATCAGCCTGGAGA-3’) and amplifications cycles consisting of an initial stage of 94°C for 3 min, 35 cycles of 95°C for 30s, 60°C for 50 s and 72°C for 1 min, and final extension at 72°C for 10 min.

For synteny analysis, the positions of *PR-1j*, *PR-1c* and *PR-1d* genes were searched in the scaffolds of genomes C-BA3, S-MG2, R-CO2, L-EC1 and L-EC2 by blastn. The scaffolds were then excised 5000bp upstream and 5000bp downstream from the starting position of *PR-1j* in the scaffolds. The resulting 10000 bp excised scaffolds were used for synteny analysis with Mummer (v. 4.0.0beta2) (Kurtz et al., 2004), using the C-BA3 sequence as the reference.

### *MpPR-1* expression data

*MpPR-1* expression data in RPKM (Reads Per Kilobase per Million mapped reads) values from the C-biotype of *M. perniciosa* in seven biological conditions (dikaryotic mycelium 14 days, basidiomata, germinating spores, green broom, initial necrosis, advanced necrosis, dry broom) were downloaded from the Witches’ Broom Disease Transcriptome Atlas (v. 1.1) (http://bioinfo08.ibi.unicamp.br/atlas/).

*MpPR-1* expression data of *M. perniciosa* treated with plant antifungal compounds were obtained from RNA-seq data. The C-BA1a isolate’s necrotrophic mycelia was initially inoculated in 100 mL of liquid MYEA media and cultivated for 5 days under agitation of 150 rpm at 30°C, then 5 mL of this initial cultivation were transferred to 50 mL of fresh MYEA liquid media containing eugenol (500μM), α-tomatin (80μM) or DMSO (250 μL, solvent control) and cultivated again under agitation of 150 rpm at 30°C for 7 days. The total RNA was extracted using the Rneasy^®^ Plant Mini Kit (Quiagen, USA) and quantified on a fluorimeter (Qubit, Invitrogen). cDNA libraries were prepared in five biological replicates for each treatment, plus biological control. The cDNA libraries were built from 1000 ng of total RNA using Illumina’s TruSeq RNA Sample Prep kit, as recommended by the manufacturer. The libraries were prepared according to Illumina’s standard procedure and sequenced on Illumina’s HiSeq 2500 sequencer. The quality of raw sequences was assessed with FastQC (v.0.11.7). Read quantification was performed by mapping the generated reads against 16084 gene models of the C-BA1a genome using Salmon (v.0.14.1) in mapping-based mode (Patro et al., 2017). Read counts were normalized to Transcripts Per Million (TPM) values for plotting. Differential expression analysis was performed with the DESeq2 (v.1.22.2) package using Wald test and Log fold change shrinkage by the *apeglm* method (IfcThreshold=0.1, s-value <0.005) (Love et al., 2014). TPM values and DESeq2 results for *MpPR-1* genes in these experimental conditions are available at Supplementary File 2.

*MpPR-1* expression data in TPM for the S-biotype was obtained from RNA-seq libraries of infected MicroTom tomato plants in 7 different time points after inoculation (12h, 24h, 48h, 5 days, 10 days, 20 days, 30 days) (Costa et al., under review, Costa, 2017). The quality of raw sequences was assessed with FastQC (v. 0.11.7). Next, Trimmomatic (v.0.36) (Bolger et al., 2014) was used to remove adaptor-containing and low-quality sequences. Quality-filtered reads were then aligned against the S-MG1 or S-MG2 reference genome using HISAT2 (v.2.1.0) with default parameters (Kim et al., 2019). Reads that mapped to coding sequences were counted with featureCounts (v.1.6.3) (Liao et al., 2014). TPM values for *MpPR-1* genes in these experimental conditions are available at Supplementary File 3.

*MrPR-1* expression data in TPM was obtained from RNA-Seq reads of *M. roreri* in the biotrophic (30 days after infection) and necrotrophic (60 days after infection) stages of frosty pod rot from (Meinhardt et al., 2014). Reads were mapped and quantified with Salmon (v.0.14.1) (Patro et al., 2017) using 17910 gene models of *M. roreri* MCA 2997 (GCA_000488995) available at Ensembl Fungi.

## Supplementary Files

**Supplementary File 1. List of fungal genomes and their source (collection site or reference publication.** This table contains the species names and biotypes of the fungal genomes used for the identification of *PR-1*-like genes, the identification names we used for the genomes, and their source, which for *M. perniciosa* and *M. roreri* isolates corresponds to their collection site, and for the other Agaricales species corresponds to their reference publication.

**Supplementary File 2.** *MpPR-1* **quantification data and differential expression results of** *M. perniciosa* **treated with eugenol or alpha-tomatin treatment.** First tab contains a matrix of expression values in Transcripts Per Million (TPM) for *MpPR-1* genes of the necrotrophic mycelia of *M. perniciosa* (C-biotype) treated with eugenol (500 μM), α-tomatin (80 μM) or DMSO (250 μL) (solvent control) for 7 days. Quantification was performed from RNA-Seq reads using the C-BA1a genome as reference. Second tab contains the results table for *MpPR-1* genes in the differential expression analysis comparing the expression profiles between Eugenol vs Control treaments, while comparison between α-tomatin vs Control treatments is shown in the third tab.

**Supplementary File 3. *MpPR-1* expression data in S-biotype infection in tomato.** Matrix of expression values in Transcripts Per Million (TPM) for *MpPR-1* genes of *M. perniciosa* S-biotype infection in MicroTom tomato in various time points of infection (12h, 24h, 48h, 5d, 10d, 20d, 30d). Quantification was performed from RNA-Seq reads using the S-MG1 (data in first tab) or S-MG2 genome (data in second tab) as reference.

**Supplementary Figure 1. Phylogenetic reconstruction of PR-1 proteins in** *Moniliophthora* **isolates (version with non-collapsed branches).** Phylogenetic relationships of PR-1 proteins identified from genomes of 18 *M. perniciosa* and 4 *M. roreri* isolates were inferred by maximum likelihood and branch support was obtained using 1000 bootstraps. The PRY1 protein of *Saccharomyces cerevisiae* was used as an outgroup.

## Acknowledgements

We are thankful to Msc. Bárbara A. Pires and Dr. Mario O. Barsottini for helping with the experiments with *M. perniciosa* treated with antifungal compounds and Msc. Leandro C. do Nascimento for performing the assembly of *Moniliophthora* genomes.

## Author contributions

J.J. and R.M.B. conceived and supervised this project. A.A.V. performed identification of PR-1-like candidate genes, evolutionary analysis, and most expression analysis from RNA seq data, executed PCR experiments and generated figures. P.J.P.L.T. and D.P.T.T. conceived the project of genomics of *Moniliophthora* isolates. P.J.P.L.T., D.P.T.T., P.F.V.P. and G.L.F. executed genomic data acquisition of *Moniliophthora* isolates. J. L.C. executed RNA-seq data acquisition of MT plants infected with S-biotype, and P.J.P.L.T. analyzed this data. P.M.T.F. performed gene prediction, annotation, and assignment of orthogroups from genomes. A. P. C. helped with genomic and RNA-seq analysis. R.M.B. conceived and executed RNA-seq data acquisition of M. *perniciosa* treated with antifungal compounds and A.A.V. analyzed this data. A.A.V. and J.J. wrote the original draft. J.J. and A.P.C. improved the design of figures. J.J., R.M.B, P.J.P.L.T, D.P.T.T., G.L.F., A.P.C., P.F.V.P. and G.A.G.P. reviewed and edited the draft. M.F.C., A.F. and G.A.G.P. contributed with project supervision and funding acquisition. All authors approved the final manuscript.

## References

Aime, M. C., & Phillips-Mora, W. (2005). The causal agents of witches’ broom and frosty pod rot of cacao (chocolate, Theobroma cacao) form a new lineage of Marasmiaceae. Mycologia, 97(5), 1012–1022. https://doi.org/10.3852/mycologia.97.5.1012

Asojo, O. A., Goud, G., Dhar, K., Loukas, A., Zhan, B., Deumic, V., Liu, S., Borgstahl, G. E. O., & Hotez, P. J. (2005). X-ray structure of Na-ASP-2, a pathogenesis-related-1 protein from the nematode parasite, Necator americanus, and a vaccine antigen for human hookworm infection. Journal of Molecular Biology, 346(3), 801–814. https://doi.org/10.1016/j.jmb.2004.12.023

Baroni, R. M., Luo, Z., Darwiche, R., Hudspeth, E. M., Schneiter, R., Pereira, G. A. G., Mondego, J. M. C., & Asojo, O. A. (2017). Crystal Structure of MpPR-1i, a SCP/TAPS protein from Moniliophthora perniciosa, the fungus that causes Witches’ Broom Disease of Cacao. Scientific Reports, 7(1), 7818. https://doi.org/10.1038/s41598-017-07887-1

Bolger, A. M., Lohse, M., & Usadel, B. (2014). Trimmomatic: A flexible trimmer for Illumina sequence data. Bioinformatics, 30(15), 2114–2120. https://doi.org/10.1093/bioinformatics/btu170

Braun, B. R., Head, W. S., Wang, M. X., & Johnson, A. D. (2000). Identification and characterization of TUP1-regulated genes in Candida albicans. Genetics, 156(1), 31–44.

Butler, G., Rasmussen, M. D., Lin, M. F., Santos, M. A. S., Sakthikumar, S., Munro, C. A., Rheinbay, E., Grabherr, M., Forche, A., Reedy, J. L., Agrafioti, I., Arnaud, M. B., Bates, S., Brown, A. J. P., Brunke, S., Costanzo, M. C., Fitzpatrick, D. A., de Groot, P. W. J., Harris, D., … Cuomo, C. A. (2009). Evolution of pathogenicity and sexual reproduction in eight Candida genomes. Nature, 459(7247), 657–662. https://doi.org/10.1038/nature08064

Camacho, C., Coulouris, G., Avagyan, V., Ma, N., Papadopoulos, J., Bealer, K., & Madden, T. L. (2009). BLAST+: Architecture and applications. BMC Bioinformatics, 10(1), 421. https://doi.org/10.1186/1471-2105-10-421

Cantacessi, C., Campbell, B. E., Visser, A., Geldhof, P., Nolan, M. J., Nisbet, A. J., Matthews,J. B., Loukas, A., Hofmann, A., Otranto, D., Sternberg, P. W., & Gasser, R. B. (2009). A portrait of the “SCP/TAPS” proteins of eukaryotes—Developing a framework for fundamental research and biotechnological outcomes. Biotechnology Advances, 27(4), 376–388. https://doi.org/10.1016/j.biotechadv.2009.02.005

Capella-Gutiérrez, S., Silla-Martínez, J. M., & Gabaldón, T. (2009). trimAl: A tool for automated alignment trimming in large-scale phylogenetic analyses. Bioinformatics (Oxford, England), 25(15), 1972–1973. https://doi.org/10.1093/bioinformatics/btp348

Chalmers, I. W., McArdle, A. J., Coulson, R. M., Wagner, M. A., Schmid, R., Hirai, H., & Hoffmann, K. F. (2008). Developmentally regulated expression, alternative splicing and distinct sub-groupings in members of the Schistosoma mansoni venom allergen-like (SmVAL) gene family. BMC Genomics, 9(1), 89. https://doi.org/10.1186/1471-2164-9-89

Chen, Y.-L., Lee, C.-Y., Cheng, K.-T., Chang, W.-H., Huang, R.-N., Nam, H. G., & Chen, Y.-R. (2014). Quantitative Peptidomics Study Reveals That a Wound-Induced Peptide from PR-1 Regulates Immune Signaling in Tomato. The Plant Cell, 26(10), 4135–4148. https://doi.org/10.1105/tpc.114.131185

Chien, P.-S., Nam, H. G., & Chen, Y.-R. (2015). A salt-regulated peptide derived from the CAP superfamily protein negatively regulates salt-stress tolerance in Arabidopsis. Journal of Experimental Botany, 66(17), 5301–5313. https://doi.org/10.1093/jxb/erv263

Choudhary, V., & Schneiter, R. (2012). Pathogen-Related Yeast (PRY) proteins and members of the CAP superfamily are secreted sterol-binding proteins. Proceedings of the National Academy of Sciences of the United States of America, 109(42), 16882–16887. https://doi.org/10.1073/pnas.1209086109

Costa, J. L. (2017). Patogenicidade e regulação hormonal na interação Moniliophthora perniciosa x Solanum lycopersicum [Tese, Centro de Energia Nuclear na Agricultura, Universidade de São Paulo]. https://doi.org/10.11606/T.64.2018.tde-04052018-100645

Darwiche, R., El Atab, O., Baroni, R. M., Teixeira, P. J. P. L., Mondego, J. M. C., Pereira, G. A. G., & Schneiter, R. (2017). Plant pathogenesis-related proteins of the cacao fungal pathogen Moniliophthora perniciosa differ in their lipid-binding specificities. The Journal of Biological Chemistry, 292(50), 20558–20569. https://doi.org/10.1074/jbc.M117.811398

Darwiche, R., Mène-Saffrané, L., Gfeller, D., Asojo, O. A., & Schneiter, R. (2017). The pathogen-related yeast protein Pry1, a member of the CAP protein superfamily, is a fatty acid-binding protein. The Journal of Biological Chemistry, 292(20), 8304–8314. https://doi.org/10.1074/jbc.M117.781880

Darwiche, R., & Schneiter, R. (2016). Cholesterol-Binding by the Yeast CAP Family Member Pry1 Requires the Presence of an Aliphatic Side Chain on Cholesterol. Journal of Steroids & Hormonal Science, 7(2). https://doi.org/10.4172/2157-7536.1000172

Deganello, J., Leal, G. A., Rossi, M. L., Peres, L. E. P., & Figueira, A. (2014). Interaction of Moniliophthora perniciosa biotypes with Micro-Tom tomato: A model system to investigate the witches’ broom disease of Theobroma cacao. Plant Pathology, 63(6), 1251–1263. https://doi.org/10.1111/ppa.12206

Ding, X., Shields, J., Allen, R., & Hussey, R. S. (2000). Molecular cloning and characterisation of a venom allergen AG5-like cDNA from Meloidogyne incognita. International Journal for Parasitology, 30(1), 77–81. https://doi.org/10.1016/s0020-7519(99)00165-4

Eddy, S. R. (2011). Accelerated Profile HMM Searches. PLoS Computational Biology, 7(10), e1002195. https://doi.org/10.1371/journal.pcbi.1002195

Emms, D. M., & Kelly, S. (2015). OrthoFinder: Solving fundamental biases in whole genome comparisons dramatically improves orthogroup inference accuracy. Genome Biology, 16(1), 157. https://doi.org/10.1186/s13059-015-0721-2

Evans, H. C. (1978). Witches’ broom disease of cocoa Crinipellis perniciosa) in Ecuador. Annals of Applied Biology, 89(2), 185–192. https://doi.org/10.1111/j.1744-7348.1978.tb07689.x

Evans, Harry C. (2007). Cacao Diseases—The Trilogy Revisited. Phytopathology^®^, 97(12), 1640–1643. https://doi.org/10.1094/PHYTO-97-12-1640

Fedorova, N. D., Khaldi, N., Joardar, V. S., Maiti, R., Amedeo, P., Anderson, M. J., Crabtree, J., Silva, J. C., Badger, J. H., Albarraq, A., Angiuoli, S., Bussey, H., Bowyer, P., Cotty, P. J., Dyer, P. S., Egan, A., Galens, K., Fraser-Liggett, C. M., Haas, B. J., … Nierman, W. C. (2008). Genomic Islands in the Pathogenic Filamentous Fungus Aspergillus fumigatus. PLOS Genetics, 4(4), e1000046. https://doi.org/10.1371/journal.pgen.1000046

Filho Tokimatu, P. M. (2018). Estudo da evolução da fitopatogenicidade em Moniliophthora perniciosa através de análises de genômica comparativa de seus biótipos [Universidade Estadual de Campinas]. http://repositorio.unicamp.br/jspui/handle/REPOSIP/334998

Fiorin, G. L., Sanchéz-Vallet, A., Thomazella, D. P. de T., do Prado, P. F. V., do Nascimento, L. C., Figueira, A. V. de O., Thomma, B. P. H. J., Pereira, G. A. G., & Teixeira, P. J. P. L. (2018). Suppression of Plant Immunity by Fungal Chitinase-like Effectors. Current Biology: CB, 28(18), 3023–3030.e5. https://doi.org/10.1016/j.cub.2018.07.055

Gamir, J., Darwiche, R., Van’t Hof, P., Choudhary, V., Stumpe, M., Schneiter, R., & Mauch, F. (2017). The sterol-binding activity of PATHOGENESIS-RELATED PROTEIN 1 reveals the mode of action of an antimicrobial protein. The Plant Journal: For Cell and Molecular Biology, 89(3), 502–509. https://doi.org/10.1111/tpj.13398

Gao, B., Allen, R., Maier, T., Davis, E. L., Baum, T. J., & Hussey, R. S. (2001). Molecular characterisation and expression of two venom allergen-like protein genes in Heterodera glycines. International Journal for Parasitology, 31(14), 1617–1625. https://doi.org/10.1016/s0020-7519(01)00300-9

Gibbs, G. M., Roelants, K., & O’Bryan, M. K. (2008). The CAP superfamily: Cysteine-rich secretory proteins, antigen 5, and pathogenesis-related 1 proteins--roles in reproduction, cancer, and immune defense. Endocrine Reviews, 29(7), 865–897. https://doi.org/10.1210/er.2008-0032

Goldman, N., & Yang, Z. (1994). A codon-based model of nucleotide substitution for protein-coding DNA sequences. Molecular Biology and Evolution, 11(5), 725–736. https://doi.org/10.1093/oxfordjournals.molbev.a040153

Griffith, G. W., & Hedger, J. N. (1994). Spatial distribution of mycelia of the liana (L-) biotype of the agaric Crinipellis perniciosa (Stahel) Singer in tropical forest. New Phytologist, 127(2), 243–259. https://doi.org/10.1111/j.1469-8137.1994.tb04276.x

Grigoriev, I. V., Nikitin, R., Haridas, S., Kuo, A., Ohm, R., Otillar, R., Riley, R., Salamov, A., Zhao, X., Korzeniewski, F., Smirnova, T., Nordberg, H., Dubchak, I., & Shabalov, I. (2014). MycoCosm portal: Gearing up for 1000 fungal genomes. Nucleic Acids Research, 42(Database issue), D699–704. https://doi.org/10.1093/nar/gkt1183

Hawdon, J. M., Narasimhan, S., & Hotez, P. J. (1999). Ancylostoma secreted protein 2: Cloning and characterization of a second member of a family of nematode secreted proteins from Ancylostoma caninum. Molecular and Biochemical Parasitology, 99(2), 149–165. https://doi.org/10.1016/s0166-6851(99)00011-0

Huerta-Cepas, J., Serra, F., & Bork, P. (2016). ETE 3: Reconstruction, Analysis, and Visualization of Phylogenomic Data. Molecular Biology and Evolution, 33(6), 1635–1638. https://doi.org/10.1093/molbev/msw046

Hunter, S., Apweiler, R., Attwood, T. K., Bairoch, A., Bateman, A., Binns, D., Bork, P., Das, U., Daugherty, L., Duquenne, L., Finn, R. D., Gough, J., Haft, D., Hulo, N., Kahn, D., Kelly, E., Laugraud, A., Letunic, I., Lonsdale, D., … Yeats, C. (2009). InterPro: The integrative protein signature database. Nucleic Acids Research, 37(Database issue), D211–D215. https://doi.org/10.1093/nar/gkn785

Kasparovsky, T., Milat, M.-L., Humbert, C., Blein, J.-P., Havel, L., & Mikes, V. (2003). Elicitation of tobacco cells with ergosterol activates a signal pathway including mobilization of internal calcium. Plant Physiology and Biochemistry, 41(5), 495–501. https://doi.org/10.1016/S0981-9428(03)00058-5

Katoh, K., & Standley, D. M. (2013). MAFFT Multiple Sequence Alignment Software Version 7: Improvements in Performance and Usability. Molecular Biology and Evolution, 30(4), 772–780. https://doi.org/10.1093/molbev/mst010

Kelleher, A., Darwiche, R., Rezende, W. C., Farias, L. P., Leite, L. C. C., Schneiter, R., & Asojo, O. A. (2014). Schistosoma mansoni venom allergen-like protein 4 (SmVAL4) is a novel lipid-binding SCP/TAPS protein that lacks the prototypical CAP motifs. Acta Crystallographica Section D: Biological Crystallography, 70(Pt 8), 2186–2196. https://doi.org/10.1107/S1399004714013315

Kerekes, J., Desjardin, D., & Desjardin, D. (2009). A monograph of the genera Crinipellis and Moniliophthora from Southeast Asia including a molecular phylogeny of the nrITS region. Fungal Diversity, 37, 101–152.

Khoza, T. G., Dubery, I. A., & Piater, L. A. (2019). Identification of Candidate Ergosterol-Responsive Proteins Associated with the Plasma Membrane of Arabidopsis thaliana. International Journal of Molecular Sciences, 20(6). https://doi.org/10.3390/ijms20061302

Kim, D., Paggi, J. M., Park, C., Bennett, C., & Salzberg, S. L. (2019). Graph-based genome alignment and genotyping with HISAT2 and HISAT-genotype. Nature Biotechnology, 37(8), 907–915. https://doi.org/10.1038/s41587-019-0201-4

Klemptner, R. L., Sherwood, J. S., Tugizimana, F., Dubery, I. A., & Piater, L. A. (2014). Ergosterol, an orphan fungal microbe-associated molecular pattern (MAMP). Molecular Plant Pathology, 15(7), 747–761. https://doi.org/10.1111/mpp.12127

Kropp, B. R., & Albee-Scott, S. (2012). Moniliophthora aurantiaca sp. Nov., a Polynesian species occurring in littoral forests. Mycotaxon, 120(1), 493–503. https://doi.org/10.5248/120.493

Kurtz, S., Phillippy, A., Delcher, A. L., Smoot, M., Shumway, M., Antonescu, C., & Salzberg, S. L. (2004). Versatile and open software for comparing large genomes. Genome Biology, 5(2), R12. https://doi.org/10.1186/gb-2004-5-2-r12

Lemoine, F., Domelevo Entfellner, J.-B., Wilkinson, E., Correia, D., Dávila Felipe, M., De Oliveira, T., & Gascuel, O. (2018). Renewing Felsenstein’s phylogenetic bootstrap in the era of big data. Nature, 556(7702), 452–456. https://doi.org/10.1038/s41586-018-0043-0

Liao, Y., Smyth, G. K., & Shi, W. (2014). featureCounts: An efficient general purpose program for assigning sequence reads to genomic features. Bioinformatics, 30(7), 923–930. https://doi.org/10.1093/bioinformatics/btt656

Lochman, J., & Mikes, V. (2006). Ergosterol treatment leads to the expression of a specific set of defence-related genes in tobacco. Plant Molecular Biology, 62(1–2), 43–51. https://doi.org/10.1007/s11103-006-9002-5

Love, M. I., Huber, W., & Anders, S. (2014). Moderated estimation of fold change and dispersion for RNA-seq data with DESeq2. Genome Biology, 15(12). https://doi.org/10.1186/s13059-014-0550-8

Lozano-Torres, J. L., Wilbers, R. H. P., Warmerdam, S., Finkers-Tomczak, A., Diaz-Granados, A., van Schaik, C. C., Helder, J., Bakker, J., Goverse, A., Schots, A., & Smant, G. (2014). Apoplastic venom allergen-like proteins of cyst nematodes modulate the activation of basal plant innate immunity by cell surface receptors. PLoS Pathogens, 10(12), e1004569. https://doi.org/10.1371/journal.ppat.1004569

Manel, S., Perrier, C., Pratlong, M., Abi-Rached, L., Paganini, J., Pontarotti, P., & Aurelle, D. (2016). Genomic resources and their influence on the detection of the signal of positive selection in genome scans. Molecular Ecology, 25(1), 170–184. https://doi.org/10.1111/mec.13468

Meinhardt, L. W., Costa, G. G. L., Thomazella, D. P., Teixeira, P. J. P., Carazzolle, M. F., Schuster, S. C., Carlson, J. E., Guiltinan, M. J., Mieczkowski, P., Farmer, A., Ramaraj, T., Crozier, J., Davis, R. E., Shao, J., Melnick, R. L., Pereira, G. A., & Bailey, B. A. (2014). Genome and secretome analysis of the hemibiotrophic fungal pathogen, Moniliophthora roreri, which causes frosty pod rot disease of cacao: Mechanisms of the biotrophic and necrotrophic phases. BMC Genomics, 15(1), 164. https://doi.org/10.1186/1471-2164-15-164

Mohd As’wad, A. W., Sariah, M., Paterson, R. R. M., Zainal Abidin, M. A., & Lima, N. (2011). Ergosterol analyses of oil palm seedlings and plants infected with Ganoderma. Crop Protection, 30(11), 1438–1442. https://doi.org/10.1016/j.cropro.2011.07.004

Nguyen, L.-T., Schmidt, H. A., von Haeseler, A., & Minh, B. Q. (2015). IQ-TREE: A fast and effective stochastic algorithm for estimating maximum-likelihood phylogenies. Molecular Biology and Evolution, 32(1), 268–274. https://doi.org/10.1093/molbev/msu300

Niveiro, N., Ramírez, N. A., Michlig, A., Lodge, D. J., & Aime, M. C. (2020). Studies of Neotropical tree pathogens in Moniliophthora: A new species M. mayarum, and new combinations for Crinipellis ticoi and C. brasiliensis. MycoKeys, 66, 39–54. https://doi.org/10.3897/mycokeys.66.48711

Nürnberger, T., Brunner, F., Kemmerling, B., & Piater, L. (2004). Innate immunity in plants and animals: Striking similarities and obvious differences. Immunological Reviews, 198, 249–266. https://doi.org/10.1111/j.0105-2896.2004.0119.x

Oleksyk, T. K., Smith, M. W., & O’Brien, S. J. (2010). Genome-wide scans for footprints of natural selection. Philosophical Transactions of the Royal Society B: Biological Sciences, 365(1537), 185–205. https://doi.org/10.1098/rstb.2009.0219

Patro, R., Duggal, G., Love, M. I., Irizarry, R. A., & Kingsford, C. (2017). Salmon provides fast and bias-aware quantification of transcript expression. Nature Methods, 14(4), 417–419. https://doi.org/10.1038/nmeth.4197

Prados-Rosales, R. C., Roldán-Rodríguez, R., Serena, C., López-Berges, M. S., Guarro, J., Martínez-del-Pozo, Á., & Di Pietro, A. (2012). A PR-1-like Protein of Fusarium oxysporum Functions in Virulence on Mammalian Hosts. The Journal of Biological Chemistry, 287(26), 21970–21979. https://doi.org/10.1074/jbc.M112.364034

Purdy, L., & Schmidt, R. (1996). STATUS OF CACAO WITCHES’ BROOM: Biology, Epidemiology, and Management. Annual Review of Phytopathology, 34(1), 573–594. https://doi.org/10.1146/annurev.phyto.34.1.573

Ranwez, V., Douzery, E. J. P., Cambon, C., Chantret, N., & Delsuc, F. (2018). MACSE v2: Toolkit for the Alignment of Coding Sequences Accounting for Frameshifts and Stop Codons. Molecular Biology and Evolution, 35(10), 2582–2584. https://doi.org/10.1093/molbev/msy159

Sambrook, J., & Russell, D. W. (2006). Purification of nucleic acids by extraction with phenol:chloroform. CSH Protocols, 2006(1). https://doi.org/10.1101/pdb.prot4455

Schneiter, R., & Di Pietro, A. (2013). The CAP protein superfamily: Function in sterol export and fungal virulence. Biomolecular Concepts, 4(5), 519–525. https://doi.org/10.1515/bmc-2013-0021

Takahashi, H. (2002). Four new species of Crinipellis and Marasmius in eastern Honshu, Japan. Mycoscience, 43(4), 343–350. https://doi.org/10.1007/S102670200050

Teixeira, Paulo J. P. L., Thomazella, D. P. T., Vidal, R. O., do Prado, P. F. V., Reis, O., Baroni, R. M., Franco, S. F., Mieczkowski, P., Pereira, G. A. G., & Mondego, J. M. C. (2012). The fungal pathogen Moniliophthora perniciosa has genes similar to plant PR-1 that are highly expressed during its interaction with cacao. PloS One, 7(9), e45929. https://doi.org/10.1371/journal.pone.0045929

Teixeira, Paulo José Pereira Lima, Costa, G. G. L., Fiorin, G. L., Pereira, G. A. G., & Mondego, J. M. C. (2013). Novel receptor-like kinases in cacao contain PR-1 extracellular domains. Molecular Plant Pathology, 14(6), 602–609. https://doi.org/10.1111/mpp.12028

Teixeira, Paulo José Pereira Lima, Thomazella, D. P. de T., & Pereira, G. A. G. (2015). Time for Chocolate: Current Understanding and New Perspectives on Cacao Witches’ Broom Disease Research. PLoS Pathogens, 11(10). https://doi.org/10.1371/journal.ppat.1005130

Teixeira, Paulo José Pereira Lima, Thomazella, D. P. de T., Reis, O., Prado, P. F. V. do, Rio, M. C. S. do, Fiorin, G. L., José, J., Costa, G. G. L., Negri, V. A., Mondego, J. M. C., Mieczkowski, P., & Pereira, G. A. G. (2014). High-Resolution Transcript Profiling of the Atypical Biotrophic Interaction between Theobroma cacao and the Fungal Pathogen Moniliophthora perniciosa. The Plant Cell, 26(11), 4245–4269. https://doi.org/10.1105/tpc.114.130807

van Loon, L. C., Rep, M., & Pieterse, C. M. J. (2006). Significance of inducible defense-related proteins in infected plants. Annual Review of Phytopathology, 44, 135–162. https://doi.org/10.1146/annurev.phyto.44.070505.143425

Yang, Z. (2007). PAML 4: Phylogenetic analysis by maximum likelihood. Molecular Biology and Evolution, 24(8), 1586–1591. https://doi.org/10.1093/molbev/msm088

Zhan, B., Liu, Y., Badamchian, M., Williamson, A., Feng, J., Loukas, A., Hawdon, J. M., & Hotez, P. J. (2003). Molecular characterisation of the Ancylostoma-secreted protein family from the adult stage of Ancylostoma caninum. International Journal for Parasitology, 33(9), 897–907. https://doi.org/10.1016/s0020-7519(03)00111-5

Zhang, J., Nielsen, R., & Yang, Z. (2005). Evaluation of an Improved Branch-Site Likelihood Method for Detecting Positive Selection at the Molecular Level. Molecular Biology and Evolution, 22(12), 2472–2479. https://doi.org/10.1093/molbev/msi237

Zhao, X. R., Lin, Q., & Brookes, P. C. (2005). Does soil ergosterol concentration provide a reliable estimate of soil fungal biomass? Soil Biology and Biochemistry, 37(2), 311–317. https://doi.org/10.1016/j.soilbio.2004.07.041

