## Supplementary Figure 1 for "Adaptive evolution of *Moniliophthora* PR-1 proteins towards its pathogenic lifestyle"

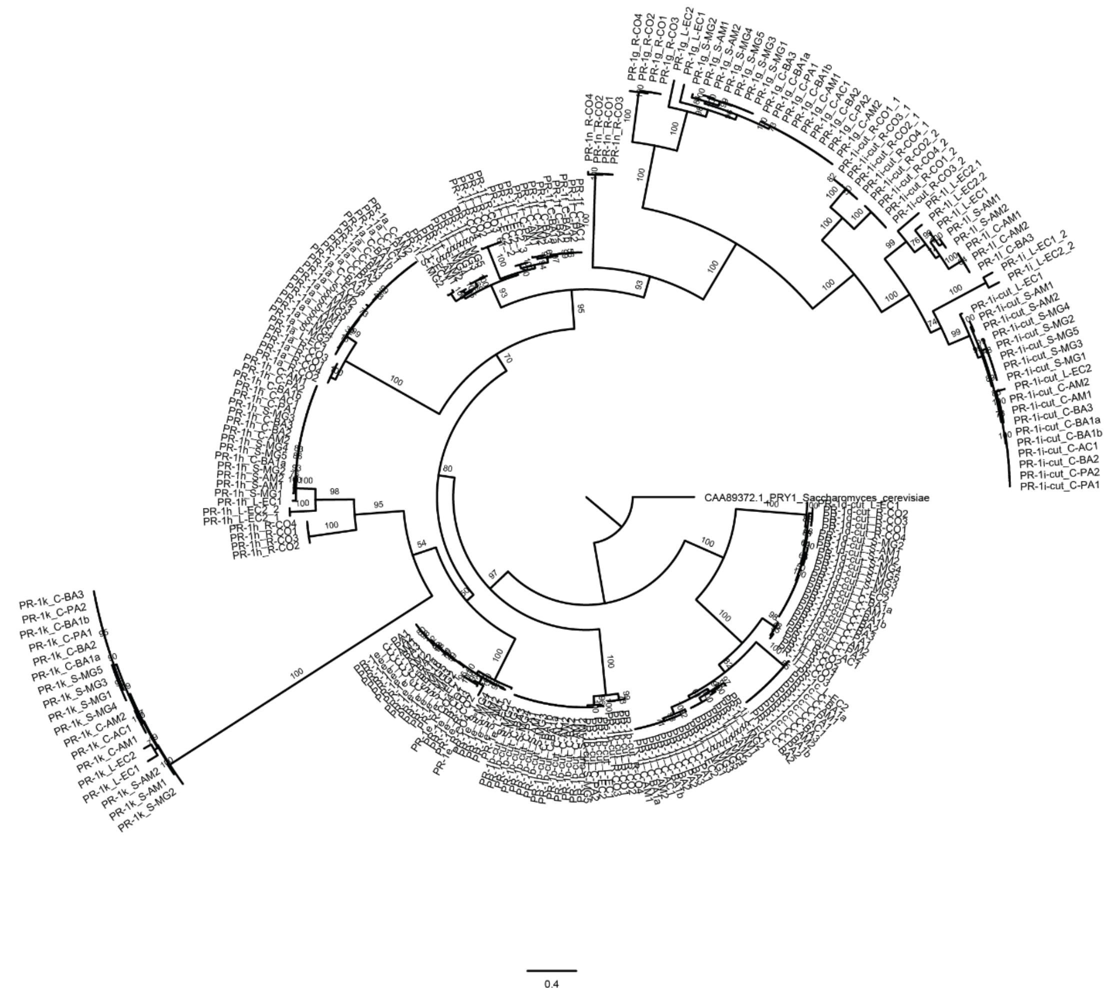


**Supplementary Figure 1. Phylogenetic reconstruction of PR-1 proteins in *Moniliophthora* isolates (version with non-collapsed branches).** Phylogenetic relationships of PR-1 proteins from 18 *M. perniciosa* and 4 M. *roreri* isolates were inferred by maximum likelihood and branch support was obtained using 1000 bootstraps. The PRY1 protein of *Saccharomyces cerevisiae* was used as an outgroup.
